# Inducible CRISPR epigenome systems mimic cocaine induced bidirectional regulation of Nab2 and Egr3

**DOI:** 10.1101/2022.09.19.508525

**Authors:** Eric Y. Choi, Daniela Franco, Catherine A. Stapf, Madeleine Gordin, Amanda Chow, Kara K. Cover, Ramesh Chandra, Mary Kay Lobo

## Abstract

Substance use disorder is a debilitating chronic disease and a leading cause of disability around the world. The nucleus accumbens (NAc) is a major brain hub that mediates reward behavior. Studies demonstrate exposure to cocaine is associated with molecular and functional imbalance in two NAc medium spiny neuron subtypes (MSNs), dopamine receptor 1 and 2 enriched D1-MSNs and D2-MSNs. Our previous reports showed that repeated cocaine exposure induced transcription factor early growth response 3 (Egr3) mRNA in NAc D1-MSNs, while reducing it in D2-MSNs. Here, we report our findings of repeated cocaine exposure inducing cell subtype specific bidirectional expression of the Egr3 corepressor NGFI-A-binding protein 2 (Nab2). Using CRISPR activation and interference (CRISPRa and CRISPRi) tools combined with Nab2 or Egr3 targeted sgRNAs, we mimicked these bidirectional changes in Neuro2a cells. Furthermore, we investigated D1-MSN and D2-MSN subtype specific expressional changes of histone lysine demethylases Kdm1a, Kdm6a and Kdm5c in NAc after repeated cocaine exposure. Since Kdm1a showed bidirectional expression patterns in D1-MSNs and D2-MSNs, like Egr3, we developed a light inducible Opto-CRISPR-KDM1a system. We were able to downregulate Egr3 and Nab2 transcripts and cause bidirectional expression changes in D1-MSNs and D2-MSNs similar to cocaine exposure in Neuro2A cells. In contrast, our Opto-CRISPR-p300 activation system induced the Egr3 and Nab2 transcripts and caused bidirectional transcription regulations in D1-MSNs and D2-MSNs. Our study sheds light on the expression patterns of Nab2 and Egr3 in specific NAc MSN subtypes in cocaine action and uses CRISPR tools to further mimic these expression patterns.

## Introduction

The nucleus accumbens (NAc) or ventral striatum, is a major brain hub that is critical for reward behaviors (Di Chiara, 2002; Wise, 2004; Russo and Nestler, 2013; Salgado and Kaplitt, 2015). Dopamine D1 receptor or D2 receptor expressing medium spiny neurons (MSNs) of the NAc project to differential outputs through the brain to constitute both the reward and ventral basal ganglia circuits (Gerfen et al., 1990; Smith et al., 2013). Previous studies demonstrate opposing roles for these two MSN subtypes in behavioral responses to psychostimulants, but the landscape of molecular changes and functional imbalance within these MSNs continue to be uncovered (Hikida et al., 2010; Lobo et al., 2010; Ferguson et al., 2011; Lobo and Nestler, 2011; Bock et al., 2013; Chandra et al., 2013; Lenz and Lobo, 2013; Engeln et al., 2020). Processes that mediate psychostimulant-induced altered transcriptional landscapes in MSN subtypes can occur through an imbalance in molecular signaling cascades, transcription factors, and epigenetic modifications of chromatin in MSNs (Lee et al., 2006; Bateup et al., 2008; Lobo et al., 2010, 2013; Lobo and Nestler, 2011; Chandra et al., 2013). This is supported by studies which demonstrate that these processes in each MSN subtype lead to differential regulation of transcription, synaptic plasticity, and behavioral responses to psychostimulants (Bateup et al., 2010; Lobo et al., 2010; Lobo and Nestler, 2011; Arango-Lievano et al., 2014; Maze et al., 2014; Chandra et al., 2015, 2017b, 2017a; Engeln et al., 2020, 2022; Cole et al., 2021).

Dysregulation of Dopamine (DA) and brain-derived neurotrophic (BDNF) signaling in the NAc are induced by repeated psychostimulant exposure resulting in altered gene expression for synaptic and behavioral plasticity geared towards repeated drug intake and seeking (Hyman et al., 2006; Russo et al., 2009; Volkow et al., 2009; McGinty et al., 2010). Here, we investigated a key molecular target of both DA and BDNF signaling, the transcription factor early growth response 3 (Egr3), and its corepressor NGFI-A binding protein 2 (Nab2) (Patwardhan et al., 1991; Yamagata et al., 1994; O’Donovan and Baraban, 1999; Jouvert et al., 2002; Roberts et al., 2006). Previous studies demonstrate that Egr3 is important for the induction of cellular programs of differentiation, proliferation, and cell death in response to environmental stimuli that rapidly and transiently induces its expression (Thiel and Cibelli, 2002; Unoki and Nakamura, 2003; Carter et al., 2007). NAB2 can modulate the activity of EGR3 by binding with EGR3 and engaging corepressor functions (Kumbrink et al., 2010). EGR3 can also bind to the Nab2 promoter and activate Nab2 transcription, thereby repressing EGR3 through a negative feedback loop (Kumbrink et al., 2005, 2010). Notably, the expression patterns of EGR3 and NAB2 have been shown to be variable depending on different cell types and stimuli (Beckmann and Wilce, 1997; Honkaniemi et al., 2000; Collins et al., 2008).

In recent years, the development of epigenetic modification tools and CRISPR tools have allowed neuroscientists to alter gene expression at specific gene loci with greater specificity compared to traditional RNAi based silencing or cDNA-based overexpression (Qi et al., 2013; Heller et al., 2014, 2016; Ran et al., 2015; Farhang et al., 2017; Klann et al., 2017; Savell et al., 2019; Duke et al., 2020; Eagle et al., 2020; Teague et al., 2022). Additionally, previous studies employed TALEs (transcription activator-like effectors) for epigenetic editing and light inducible epigenetic editing (Konermann et al., 2013). However, the complex and lengthy procedures of TALE engineering make CRISPR approaches more favorable. Here, we employ D1-Cre and D2-Cre specific RiboTag mice (Gong et al., 2007; Sanz et al., 2009, 2013; Gerfen et al., 2013; Chandra et al., 2015) to further explore Nab2 and other Egr3 targets in MS subtypes after repeated cocaine exposure and use these CRISPR tools in Neuro2A cells to explore Egr3 and Nab2 dynamics.

## Materials and Methods Mice

Wild type C57BL/6J mice were purchased from Jackson Laboratory and used for ChIP experiments. Homozygous RiboTag (RT) mice on a C57BL/6J background expressing a Cre inducible HA-Rpl22 were purchased from Jackson Laboratory. D1-Cre hemizygote (line FK150) and D2-Cre hemizygote (line ER44) BAC transgenic mice from GENSAT on C57BL/6J background were crossed with RT mice to generate D1-Cre-RT and D2-Cre-RT mice and used for cell type specific isolation of ribosome associated mRNA. All mice used for experiments were maintained on a 12-hour light/dark cycle with *libitum* access to food and water. All experiments were conducted in accordance with the guidelines set up by the Institutional Animal Care and Use Committee at the University of Maryland School of Medicine.

### Repeated Cocaine Treatment

Male D1-Cre-RT, D2-Cre-RT and C57BL/6J wild type mice received 7 daily intraperitoneal injections of cocaine (20mg/kg) or 0.9% saline in the home cage. NAc tissue punches were collected 24 hours after the last injection for molecular biology experiments or cross-linked in 1% formaldehyde for ChIP experiments. Cocaine hydrochloride (Sigma) was dissolved in sterile saline. The dose of cocaine and the 24 hour abstinent time point was selected based on previous studies showing important cocaine-mediated transcriptional and plasticity changes at the time point (Maze et al., 2010; Russo et al., 2010; Kim et al., 2011; Feng et al., 2014).

### Cell Culture Transfection

Neuro2A cell cultures were maintained in DMEM supplemented with 10% FBS and penicillin/streptomycin (all from Thermo Fisher Scientific). Cell cultures of passage number between 5-10 were used for the entire experiment. 24 hours prior to transfection, cells were seeded at the density of 1.5 × 10^5^ cells per well in 24 well plates. Transfection was performed using Invitrogen LF3000 reagents following the manufacturer’s standard protocol. 6 hours after the start of transfection, cells received fresh media change with DMEM supplemented with 10% FBS and penicillin/streptomycin. For CRISPRa and CRISPRi experiments, cells were harvested 48 hours post transfection for RNA, protein, or Cut&Run sample preparation. For Opto-CRISPRa and Opto-CRISPRi experiments, cells received light stimulation 48 hours post transfection, and harvested 1hr post light stimulation for RNA sample, 4hr post light stimulation for protein sample, or immediately for Cut&Run sample.

### Opto-CRISPR Light Stimulation

A custom built 24 well light plate apparatus with 460nm LED bulbs set to emit 1mW of light was used for light stimulation (Gerhardt et al., 2016). On the day of light stimulation (48 hours post transfection), cells plated on 24 well plates received media change to colorless BrightCell NEUMO Photostable media (EMD Millipore SCM146-1) supplemented with 1X BrightCell SOS Neuronal Supplement (EMD Millipore SCM147-1) and 1% penicillin/streptomycin at least 3 hours prior to the start of light stimulation in order to avoid blue light-induced gene alterations (Duke et al., 2019). 2 hour steady state light stimulation iris protocol was loaded onto the light plate apparatus (taborlab.github.io/iris/). The 24 well plates with transfected cells were inserted into the light plate apparatus and placed back into the cell incubator set at 37°C and 5% CO_2_ level for 2 hour light stimulation. After 2 hours of light stimulation, the cells remained in the incubator and was harvested 1hr post light stimulation for RNA samples, 4hr post light stimulation for protein samples, or immediately for Cut and Run samples.

### Western Blotting

Cell lysates were prepared with RIPA buffer (Sigma-Aldrich R0278) supplemented with Phosphatase Inhibitor Cocktail 2 (Sigma-Aldrich P5726), Phosphatase Inhibitor Cocktail 3 (Sigma-Aldrich P0044), Complete Protease Inhibitor Cocktail (Roche 11873580001).

Electrophoresis was performed using Mini-Protean TGX Gels (Biorad 4569033) followed by the transfer onto a nitrocellulose membrane (Biorad 1620094). After blocking in 5% skim milk/Tris-buffered saline supplemented with 0.1% Tween 20 (TBST), membranes were incubated with primary antibody overnight at 4°C followed by secondary antibody incubation for 1 hour at room temperature. Blot images were captured using Biorad ChemiDoc MP Imager, and the raw band intensities were quantified relative to GAPDH band intensities using ImageJ (NIH). For Opto-CRISPR-KDM1A and Opto-CRISPR-p300 western blot quantification analyses, GAPDH normalized NAB2 band intensities from 2 hours light stimulation samples were normalized to GAPDH normalized NAB2 band intensities from no light stimulation control samples.

Consistently, GAPDH normalized EGR3 band intensities from 2 hours light stimulation samples were normalized to GAPDH normalized EGR3 band intensities from no light stimulation control samples. The following primary antibodies were used at 1:1000 dilution: NAB2 (Santa Cruz Biotechnology SC-22815), EGR3 (Santa Cruz Biotechnology SC-191), GAPDH (Cell Signaling 2118s).

### Chromatin Immunoprecipitation

Fresh NAc punches were prepared for ChIP as described previously (Chandra et al., 2015). Briefly, four 14 NAc punches per animal (5 male mice pooled per sample) were collected, cross-linked with 1% formaldehyde, and quenched with 2M glycine before being flash frozen on dry ice and stored in −80°C. Before sample sonication, IgG magnetic beads (Invitrogen; Sheep anti-rabbit 11202D) were incubated in blocking solution (0.5% BSA in PBS) which contains 15ug of anti-Nab2 antibody (Santa Cruz SC22815) per reaction at 4°C overnight under constant rotation. NAc tissues were dounce homogenized in 1ml lysis buffer (50mM HEPES-KOH, pH 7.5, 140mM NaCl, 1mM EDTA, 10% glycerol, 0.5% NP-40, 0.25% Triton X-100, and protease inhibitors) and placed on rotator at 4°C for 10 min. Samples were centrifuged at 1350 X g for 5 min at 4°C and the pellet was resuspended in 1ml of lysis buffer 2 (10mM Tris-HCl, pH 8.0, 200mM NaCl, 1mM EDTA, 0.5mM EGTA, and protease inhibitors) and incubated gently on shaker at room temperature for 10 min. Then, nuclei were pelleted by centrifugation at 1350 X g for 5 min at 4°C and resuspended in 300ul of lysis buffer 3 (10mM Tris-HCl, pH 8.0, 100mM NaCl, 1mM EDTA, 0.5mM EGTA, 0.1% Na-deoxycholate, 0.5% *N*-lauroylsarcosine, and protease inhibitors). Chromatins were sheared to an average length of 500-700bp by the Diagenode Bioruptor Pico Sonicator using eight cycles of 30 seconds on and off at 4°C. Then, 1/10 volume of 10% Triton X-100 was added to the sonicated lysate to permeate the nuclear membrane. Samples were centrifuged at 20,000 X g for 10 min at 4°C to pellet the debris. After washing and resuspension of the antibody bead conjugates, the mixtures were added to each chromatin sample and incubated for 16 hours under constant rotation at 4°C. Samples were then washed on magnetic stands and reverse cross-linked at 65°C overnight. Chromatin was purified using a PCR purification kit (Qiagen 28104) and used for qPCR analysis normalized to their respective input controls as previously described. The list of primers used for ChIP-qPCR and their sequences are available in table S1.

### Cut & Run

Neuro2a cells were harvested 48 hours post transfection for CRISPRa and CRISPRi experiments, flash frozen on dry ice, and stored at −80°C. For Opto-CRISPRa and Opto-CRISPRi experiments, Neuro2a cells were harvested immediately post light stimulation, flash frozen on dry ice, and stored at −80°C. Cut & Run experiment was performed using Cell Signaling Cut & Run Kit (86652) following the manufacturer’s standard protocol. Cells were homogenized in 1ml dounce homogenizer, 20 ups and downs using pestle A and then another 20 ups and downs using pestle B, in wash buffer supplemented with spermidine and protease inhibitor cocktail according to the manufacturer’s recipe. Enriched chromatins were purified using Cell Signaling ChIP and Cut & Run DNA Purification Kit (14209) following the manufacturer’s standard protocol. The following antibody were used at 2ul per sample for negative control and 5ul per sample for all remaining antibodies: Normal Rabbit IgG (Cell Signaling 66362), H3K4me3 (Cell Signaling 9751), H3K27AC (Cell Signaling 8173), HA (Cell Signaling 3724).

### Polyribosome Immunoprecipitation and RNA Isolation

Immunoprecipitation of polyribosome was prepared from NAc of male D1-Cre-RT and D2-Cre-RT mice according to our previous study (Chandra et al., 2015). In brief, four 14-gauge NAc punches per animal (four animals pooled per sample) were collected and homogenized by douncing in homogenization buffer. 800ul of the homogenate was added directly to the HA-coupled magnetic beads (Invitrogen: 100.03D; Covance: MMS-101R) for constant rotation overnight at 4°C. The following day, beads were washed three times in high salt buffer using a magnetic stand. Finally, RNA was extracted using the MicroElute Total RNA Kit (Omega) according to the manufacturer’s standard protocols. RNA was quantified with a NanoDrop (Thermo Scientific).

### RNA Extraction and cDNA Synthesis

RNA extraction samples were prepared from Neuro2a cells using Qiagen RNeasy micro plus kit (Qiagen 74034). Neuro2a cells were harvested directly from the wells with application of Buffer RLT and physical scraping using a cell scraper. The cell lysate then underwent series of affinity column-based extraction and purification following the manufacturer’s standard protocol. Purified RNA samples were quantified, and 500ng cDNA samples were synthesized using Biorad iScript cDNA synthesis kit following the manufacturer’s standard protocol (Biorad 1708891). Synthesized cDNA samples were diluted 10 folds and stored in −20°C.

### qPCR Analysis

qRT-PCR and Cut&Run qPCR experiments were carried out with PerfeCTa SYBR Green FastMIX (Quantabio 95072-012) on Biorad CFX384 Real-Time Thermal Cycler. Gene specific primer sets were obtained from MGH PrimerBank (https://pga.mgh.harvard.edu/primerbank/) or designed using NCBI Primer BLAST (https://www.ncbi.nlm.nih.gov/tools/primer-blast/). The primer sequences used in the experiments: Nab2 Primer F – CATTCTCCATCCTTGCACTCG; Nab2 Primer R – CCCTGCCTGTTATTGCTTGA; Egr3 Primer F – CCGGTGACCATGAGCAGTT; Egr3 Primer R – TAATGGGCTACCGAGTCGC; Nab2 Cut&Run Primer F – CCGGTTCTGCCCACGCCCTC; Nab2 Cut&Run Primer R – TCCACCGCTGTCCAGGCTCC; Egr3 Cut&Run Primer F – GCTGCACGTTGGCTGCGG; Egr3 Cut&Run Primer R – CCTAGCTAGCTCACTGCTGCC; Nab2 ChIP Primer 1 F – CGCCAGGCAAAGAGCAGCGC; Nab2 ChIP Primer 1 R – CGGAGTCCAACTCCACCCATCT; Nab2 ChIP Primer 2 F – GGGGTCTAAAGGGTGGAGGTCG; Nab2 ChIP Primer 2 R – CAGGGGGCTCATGTGCCAGGT.

### Immunocytochemistry

Neuro2a cells plated on glass coverslips were fixed in 4% paraformaldehyde for 30 minutes and then washed with phosphate buffered saline (PBS). The coverslips were then placed in blocking solution (PBS supplemented with 2% normal donkey serum, 1% bovine serum albumin, 0.1% triton X-100, 0.05% Tween 20, and 0.05% sodium azide) for 1 hour at room temperature and then incubated at 4°C overnight with the following primary antibodies: chicken anti-GFP (1:5000, Aves Labs GFP-1020), mouse anti-HA tag (1:400, Abcam ab18181), rabbit anti-FLAG tag (1:400, Sigma F7425). After series of washing with PBS, the coverslips were incubated with fluorophore conjugated secondary antibodies at 1:400 dilution for either 2 hours at room temperature or at 4°C overnight. Following another series of washing with PBS, the coverslips were mounted on glass slides with DAPI containing Vectashield mounting medium (Vector laboratories H-1200). Z-stack Images were acquired with Leica SP8 confocal microscope and then processed on ImageJ (NIH).

### Constructs

Cre inducible pAAV-EF1a-DIO-dSaCas9-CIBN-HA vector and pAAV-EF1a-DIO-Cry2-KDM1A-FLAG vector were commercially made from VectorBuilder Inc. pAAV-EF1a-DIO-Cry2-p300core-FLAG vector was made by taking out KDM1A sequence out of the pAAV-EF1a-DIO-Cry2-KDM1A-FLAG vector using the restriction sites EcoRI and NotI, and cloning in p300core insert fragment into the EcoRI and NotI sites. pAAV-EF1a-DIO-dSaCas9-VP64-HA vector and pAAV-EF1a-DIO-dSaCas9-KRAB-HA were made by taking out CIBN sequence out of the pAAV-EF1a-DIO-dSaCas9-CIBN-HA vector using the restriction sites EcoRI and NotI, and cloning in VP64 or KRAB insert fragments into the EcoRI and NotI sites. sgRNA vectors with an independent Cre inducible EGFP reporter were designed using PX552 vector gifted from Feng Zhang Lab (Addgene plasmid #60958; http://n2t.net/addgene:60958; RRID:Addgene_60958). Conventional EGFP sequence in the original vector was taken out using the restriction sites SalI and EcoRI and replaced with DIO-NLS-EGFP sequence using the same SalI and EcoRI sites. The sequences of all the vector used in this study were confirmed by conventional Sanger sequencing.

### Statistical Analysis

Student *t* test and one-way ANOVA were used with *P* < 0.05 considered statistically significant. For multiple testing corrections, Tukey’s post hoc test was utilized as indicated in the results section. All the statistical analyses were performed with GraphPad Prism 6 (GraphPad Software Inc.).

## Results

### Nab2 is bidirectionally regulated in NAc D1-MSNs versus D2-MSNs after repeated exposure to cocaine

We first investigated the regulation of Nab2 expression in the NAc after seven daily injections of cocaine (20mg/kg) followed by a 24hour abstinent period, which is a critical time point for transcriptional and plasticity changes occurring in the NAc as previously reported (Fig1 A) (Maze et al., 2010; Russo et al., 2010; Kim et al., 2011; Feng et al., 2014; Chandra et al., 2015, 2017b, 2017a). Using qRT-PCR, we did not observe a statistically significant change in Nab2 mRNA expression, however, there is a trend toward an increase in Nab2 mRNA expression after repeated cocaine exposure in bulk NAc tissue (Student’s *t* test, p=0.056; mRNA n= 6-7 per group, *t*_(10)_= 2.133, p<0.05; Fig1 B). Next, we performed chromatin immunoprecipitation (ChIP) to examine hallmark activation and repressive histone methylation modifications on lysine residues, H3K9me2, H3K27me3, and H3K4me3 (Shi et al., 2004; Wysocka et al., 2006; Robison and Nestler, 2011; Ferrari et al., 2014; Nestler, 2014) on the Nab2 promoter. Interestingly, we observed that repression associated H3K9me2 and H3K27me3 modifications on Nab2 promoter were bidirectionally regulated in the NAc. H3K9me2 enrichment on Nab2 promoter was decreased while H3K27me3 enrichment on Nab2 promoter was increased in the NAc after repeated exposure to cocaine compared to saline control mice (Student’s *t* test, *p<0.05, **p<0.01; pooled ChIP sample n= 5-6 per group, *t*_(9)_= 2.309, p<0.05; pooled ChIP sample n= 5-6 per group, *t*_(9)_= 3.774, p<0.05; Fig1 C). Furthermore, we saw an increase in activation associated H3K4me3 modification on Nab2 chromatin after repeated exposure to cocaine compared to saline controls (Student’s *t* test, **p<0.01; pooled ChIP sample n= 6 per group, *t*_(10)_= 3.903, p<0.05; Fig1 C). We also examined the enrichment of histone demethylase KDM1A, which has been previously reported to induce demethylation at H3K4 and H3K9 (Shi et al., 2004; Metzger et al., 2005; Rudolph et al., 2013), on the Nab2 promoter. Interestingly, we saw an increase in the enrichment of KDM1A on the Nab2 chromatin promoter (Student’s *t* test, *p<0.05; pooled ChIP sample n= 6 per group, *t*_(10)_= 3.070, p<0.05; Fig1 C). These complex and bidirectional findings of transcriptional regulation of Nab2 in the heterogeneous NAc suggested a need for cell subtype specific analysis to elucidate Nab2 dynamics in cocaine action.

**Fig. 1.**
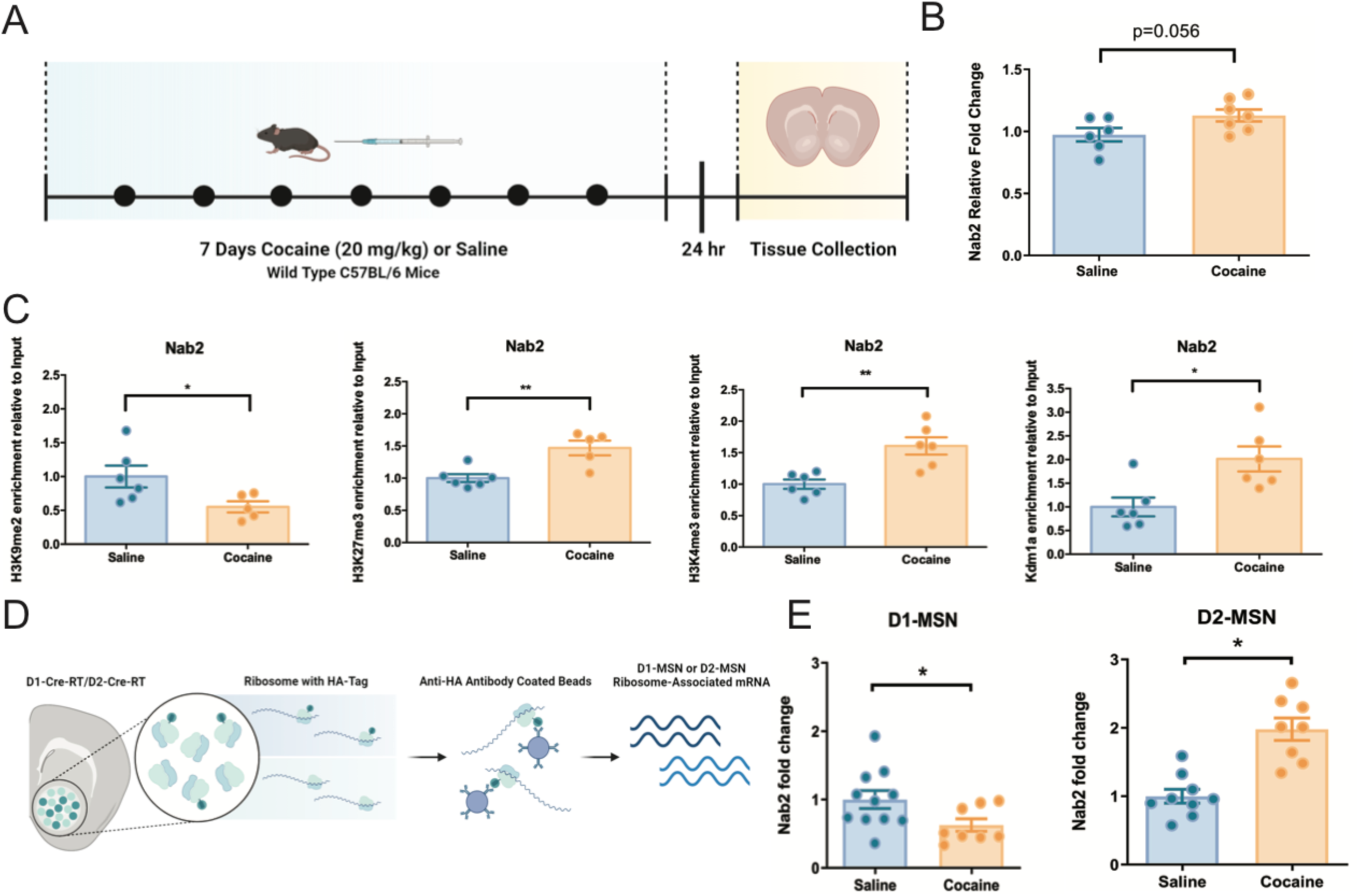
Nab2 ribosome-associated mRNA expression after repeated cocaine exposure. **A**. Male mice received intraperitoneal injection of cocaine (20mg/kg) for 7 consecutive days. NAc punches were harvested 24 hours post last cocaine injection. **B**. qRT-PCR analysis shows a trend towards an increase in Nab2 mRNA expression in the NAc after repeated cocaine exposure in male mice (saline n= 6, cocaine n= 7). **C**. Transcription repression associated H3K9me2 and H3K27me3 marks show bidirectional enrichment on Nab2 promoter after repeated exposure to cocaine. A transcription activation associated H3K4me3 mark and a histone demethylase Kdm1a both showed an increased enrichment on Nab2 promoter after repeated exposure to cocaine (H3K4me3 saline n= 6, cocaine n= 6; H3K9me2 saline n= 6, cocaine n= 5; H3K27me3 saline n= 6, cocaine n= 5; Kdm1a saline n= 6, cocaine n= 6). **D**. D1-Cre-RT and D2-Cre-RT mice allow cell subtype specific isolation of ribosome associated mRNA. **E**. Repeated cocaine exposure reduced Nab2 mRNA expression in D1-MSNs while inducing Nab2 mRNA expression in D2-MSNs (D1-MSNs saline n= 11, cocaine n= 8; D2-MSNs saline n= 9, cocaine n= 8). Each bar represents mean ± SEM. \**p*<0.05, \*\**p*<0.01.

Using a Cre inducible RiboTag mouse line crossed with D1 Cre and D2 Cre mouse lines, we generated mouses lines with D1-or D2-MSN specific expression of ribosomal subunit Rpl22 labeled with hemagglutinin (HA) protein which allow cell subtype specific polyribosome immunoprecipitation with an HA antibody (D1-Cre-RT and D2-Cre-RT). These mouse lines and similar approaches allow isolation of ribosome associated mRNA from each MSN subtypes, as previously published (Fig1 D) (Lobo et al., 2006; Heiman et al., 2008; Ena et al., 2013; Rothwell et al., 2014; Chandra et al., 2015). Seven daily injections of cocaine (20mg/kg) followed by a 24 hours of abstinent period resulted in reduced Nab2 mRNA expression in D1-MSNs while inducing it in D2-MSNs (Student’s *t* test, *p<0.05; pooled ChIP sample n= 8-11 per group, *t*_(17)_= 2.156, p<0.05; pooled ChIP sample n= 8-9 per group, *t*_(15)_= 5.195, p<0.05; Fig1 E). Notably, the mRNA expression patterns of Nab2 in D1 and D2-MSNs were in the opposite direction to our previously reported mRNA expression patterns of Egr3 (Chandra et al., 2015).

### CRISPRi and CRISPRa induced perturbation of Nab2 and Egr3 expression mimics cocaine-induced bidirectional regulation of these transcripts

We next adopted a CRISPR interference (CRISPRi) or CRISPR activation (CRISPRa) system in Neuro2a cells to determine if we can recapitulate the cocaine-induced Egr3 and Nab2 transcriptional regulation observed with repeated cocaine in NAc cell types. We transfected Neuro2a cells with Cre inducible DIO-dCas9-KRAB-HA (CRISPRi) or DIO-dCas9-VP64-HA (CRISPRa); an sgRNA targeting Egr3, Nab2, or lacZ promoters with an EGFP reporter, and Cre-recombinase (Fig2 A). This system allows the flexibility to freely exchange sgRNA vectors for different gene targets and allows targeting to Cre expressing cells. To validate the Cre-induced expression of our vectors, we performed immunocytochemistry on Neuro2a cells transfected with CRISPRi and lacZ targeting sgRNA with or without Cre recombinase vector. 48 hours post transfection, samples were fixed and immunostained with anti-GFP, anti-HA, and DAPI was labeled. We observed GFP and HA signals only in Cre positive cells (Fig2 B). In Cre negative cells, we did not observe signals from GFP and HA staining but only observed DAPI signals (Fig2 B). We performed the same experiment with the CRISPRa and lacZ targeting sgRNA with or without Cre recombinase vector. Consistent with our results using CRISPRi, we only observed GFP and HA signals from Cre positive cells (Fig2 C).

**Fig. 2.**
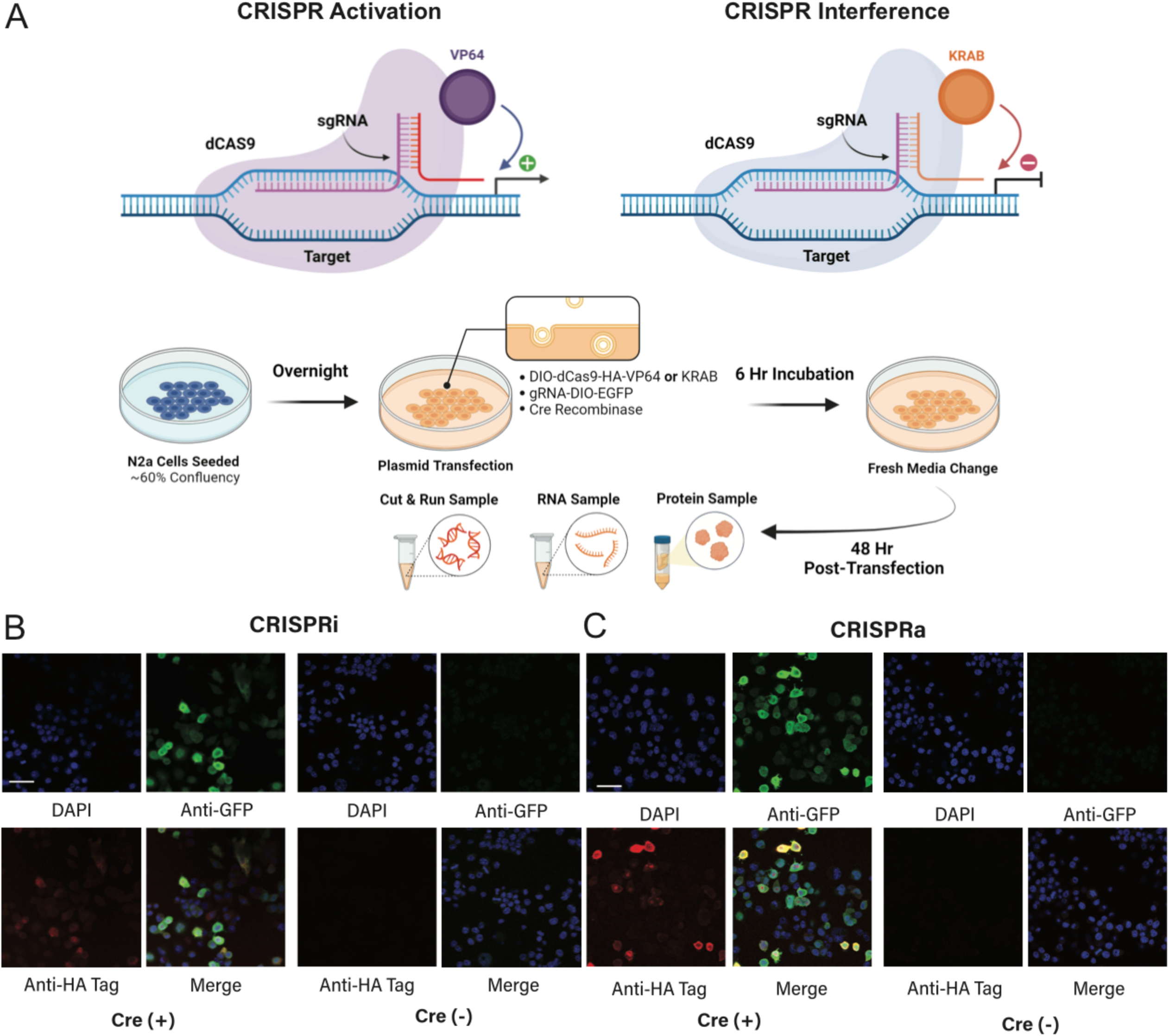
Cell subtype specific CRISPRa transcriptional activation and CRISPRi transcriptional interference. **A**. Illustration of a Cre recombinase dependent expression of a catalytically dead saCas9 fused with VP64 or KRAB in Neuro2a cell-line. Transfected cells are harvested 48 hours post transfection. **B and C**. Immunocytochemistry of Neuro2a cells transfected with CRISPRi and CRISPRa systems display a Cre inducible expression as observed by GFP (green) and HA-Tag (red) expression. DAPI signal is blue. Scale bar, 50μM.

To determine if we can use CRISPRi and CRISPRa systems to target Nab2 and Egr3, we designed sgRNAs against Nab2 and Egr3. For Nab2 sgRNAs, two ideal spacer sequences followed by NGRRN PAM sequence were identified 363bp and 729bp upstream of the Nab2 transcription start site (Ran et al., 2015). Egr3 sgRNA was designed in a similar manner with ideal spacer sequence 567bp upstream of the Egr3 transcription start site. Neuro2a cells transfected with Nab2 sgRNAs and CRISPRi displayed significant reduction of Nab2 mRNA expression while Egr3 mRNA was upregulated compared to lacZ sgRNA controls (One-way ANOVA, Nab2: interaction *F*_(3, 32)_= 30.46; p<0.0001, Tukey post-test: *p<0.05, **p<0.01, ****p<0.0001. Egr3: interaction *F*_(3, 32)_= 15.74; p<0.0001, Tukey post-test: *p<0.05, **p<0.01; mRNA n= 9 per group; Fig3 A). In cells transfected with Egr3 sgRNA and CRISPRi, we observed reduction of Egr3 mRNA and upregulation of Nab2 mRNA compared to lacZ sgRNA controls (One-way ANOVA, Nab2: interaction *F*_(3, 32)_= 30.46; p<0.01, Tukey post-test: *p<0.05, **p<0.01, ****p<0.0001. Egr3: interaction *F*_(3, 32)_= 15.74; p<0.01, Tukey post-test: *p<0.05, **p<0.01; mRNA n= 9 per group; Fig3 A). These data are consistent with the bidirectional regulation of Nab2 and Egr3 that we observed in NAc cell types after repeated cocaine exposure. Using CRISPRa with Nab2 or Egr3 sgRNAs, we successfully activated Nab2 mRNA or Egr3 mRNA with their respective sgRNAs when compared to lacZ sgRNA control (One-way ANOVA, Nab2: interaction *F*_(3, 32)_= 10.35; p<0.0001, Tukey post-test: **p<0.01. Egr3: interaction *F*_(3, 32)_= 5.848; p<0.01, Tukey post-test: **p<0.01; mRNA n= 9 per group; Fig3 B). With each sgRNA and CRISPRa, we did not observe the bidirectional regulation of the other gene that we observed with CRISPRi. To confirm these changes are consistent at the protein levels, we performed western blots with a well validated NAB2 and EGR3 antibody. Neuro2a cells transfected with Nab2 sgRNAs and CRISPRi displayed significant reduction of NAB2 protein expression while EGR3 protein expression was upregulated compared to lacZ sgRNA controls (One-way ANOVA, Nab2: interaction *F*_(3, 8)_= 27.33; p= 0.0001, Tukey post-test: *p<0.05, **p<0.01. Egr3: interaction *F*_(3, 8)_= 26.03; p<0.001, Tukey post-test: *p<0.05, **p<0.01; protein n= 3 per group; Fig3 C). In cells transfected with Egr3 sgRNA and CRISPRi, we observed reduction of EGR3 protein expression while NAB2 protein expression was upregulated compared to lacZ sgRNA controls (One-way ANOVA, Nab2: interaction *F*_(3, 8)_= 27.33; p= 0.0001, Tukey post-test: *p<0.05, **p<0.01. Egr3: interaction *F*_(3, 8)_= 26.03; p<0.001, Tukey post-test: *p<0.05, **p<0.01; protein n= 3 per group; Fig3 C). Contrarily, cells transfected with Nab2 sgRNA and CRISPRa showed significant upregulation of NAB2 protein expression, and reduced EGR3 protein expression compared to lacZ sgRNA controls (One-way ANOVA, Nab2: interaction *F*_(3, 8)_= 6.264; p= 0.05, Tukey post-test: *p<0.05. Egr3: interaction *F*_(3, 8)_= 154.6; p<0.0001, Tukey post-test: **p<0.01, ****p<0.0001; protein n= 3 per group; Fig3 C). In cells transfected with CRISPRa targeted by Egr3 sgRNA, we saw an induction of EGR3 protein expression while NAB2 protein expression did not change compared to lacz sgRNA controls (One-way ANOVA, Nab2: interaction *F*_(3, 8)_= 6.264; p<0.05, Tukey post-test: *p<0.05. Egr3: interaction *F*_(3, 8)_= 154.6; p<0.0001, Tukey post-test: **p<0.01, ****p<0.0001; protein n= 3 per group; Fig3 C). Notably, we observed that transfection with Nab2-363 gRNA was able to induce the changes in Nab2 and Egr3 in both the CRISPRi and CRISPRa systems compared to Nab2-729, suggesting that targeting closer to the ATG site provides be improved efficiency for targeting of Nab2 and we focused on this gRNA target in subsequent validation of CRISRPi/a.

**Fig. 3.**
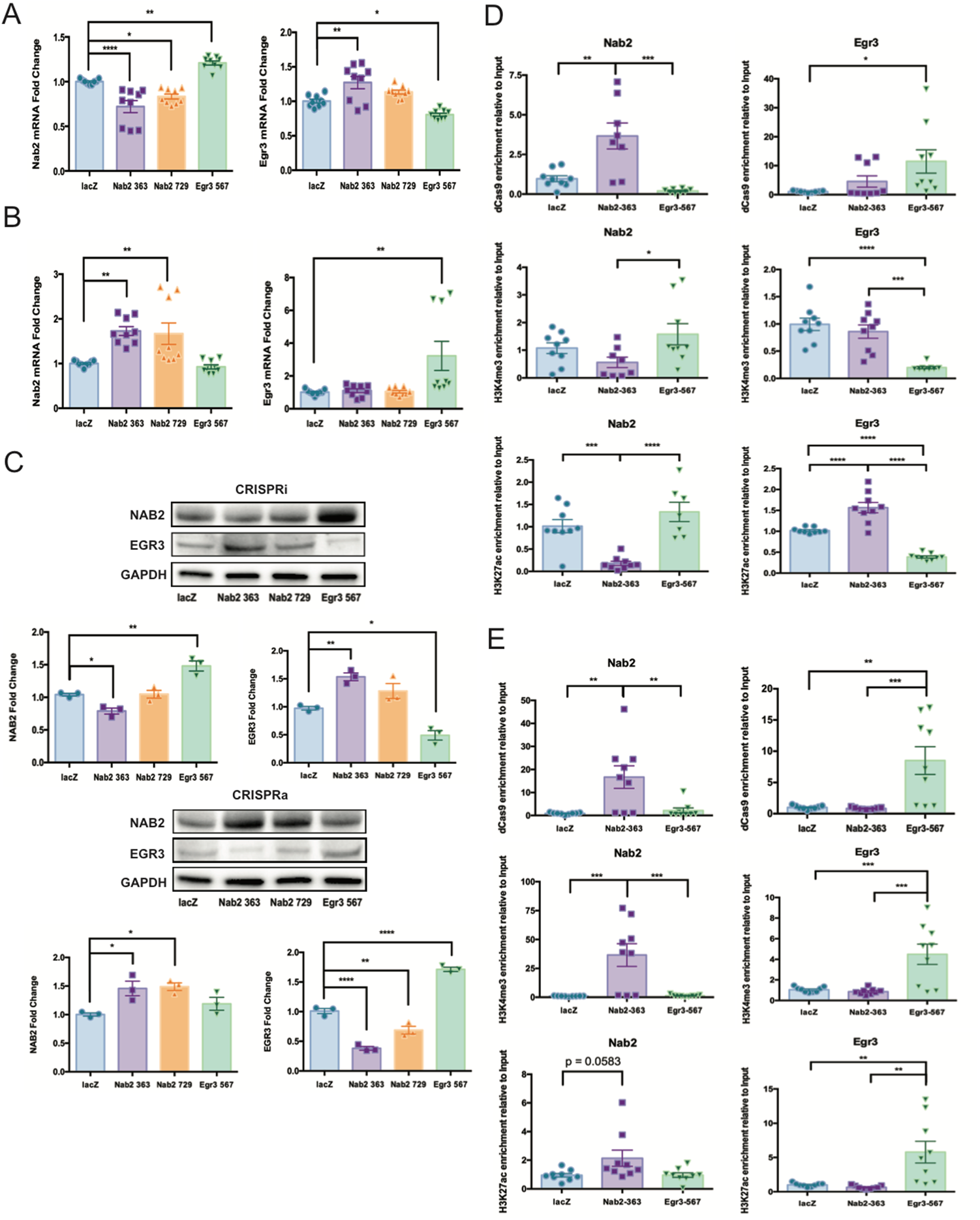
CRISPRi and CRISPRa manipulation of Nab2 and Egr3 transcription. **A**. qRT-PCR shows Nab2 targeted CRISPRi repressed Nab2 mRNA while inducing Egr3 mRNA expression, and Egr3 targeted CRISPRi repressed Egr3 mRNA while inducing Nab2 mRNA expression in Neuro2a cells (n= 9 per condition). **B**. Conversely, Nab2 targeted CRISPRa activates Nab2 mRNA expression, and Egr3 targeted CRISPRa activates Egr3 mRNA expression in Neuro2a cells (n= 9 per condition). **C**. Western blots using Neuro2a cells transfected with Nab2 targeted CRISPRi show downregulation of NAB2 and upregulation of EGR3, while cells transfected with Nab2 targeted CRISPRa show upregulation of NAB2 and downregulation of EGR3. Consistently, Neuro2a cells transfected with Egr3 targeted CRISPRi samples show downregulation of EGR3 and upregulation of NAB2, while cells transfected with Egr3 targeted CRISPRa show upregulation of EGR3 but did not change NAB2 levels (n= 3 per condition). **D**. Cut and Run of Neuro2a cells transfected with CRISPRi show HA-tag enrichment on the promoters of respective genes targeted by sgRNAs. Transcriptional activation marks, H3K4me3 and H3K27ac, display reduced enrichment on the promoters of respective genes targeted by sgRNAs compared to lacZ controls (n= 7-9 per condition). **E**. Cut and Run of Neuro2a cells transfected with CRISPRa show HA-tag, H3K4me3, and H3K27ac enrichment on the promoters of respective genes targeted by sgRNAs compared to lacZ controls (n= 9 per condition). Each bar represents mean ± SEM. \**p*<0.05, \*\**p*<0.01, \*\*\**p*<0.001, \*\*\*\**p*<0.0001.

To validate the specificity of CRISPRi and CRISPRa in our findings, we performed Cut & Run experiments (Skene and Henikoff, 2017) with anti-HA to target the HA-tag on the CRISPRi/a vectors, as well as antibodies against hallmark activation and repressive histone methylation and acetylation modifications H3K4me3 and H3K27ac (Wysocka et al., 2006; Heintzman et al., 2007, 2009; Wang et al., 2008; Creyghton et al., 2010; Rudolph et al., 2013). In Neuro2a cells transfected with Nab2 sgRNA and CRISPRi system, we saw an increase in the enrichment of CRISPRi complex on Nab2 promoter compared to lacZ sgRNA controls (One-way ANOVA, interaction *F*_(2, 22)_= 14.78; p<0.0001, Tukey post-test: **p<0.01, ***p<0.001; Cut and Run chromatin n= 7-9 per group; Fig3 D). In samples with the Egr3 sgRNA CRISPRi system, we observed an increase in the enrichment of CRISPRi complex on Egr3 promoter compared to lacZ sgRNA controls (One-way ANOVA, interaction *F*_(2, 22)_= 4.200; p= 0.0273, Tukey post-test: *p<0.05; Cut and Run chromatin n=7-9 per group; Fig3 D). Interestingly, we did not observe a significant change in the enrichment of H3K4me3 on Nab2 promoter with the Nab2 sgRNA CRISPRi system, but we did observe a decrease of H3k4me3 enrichment on Egr3 promoter with the Egr3 sgRNA CRISPRi system (One-way ANOVA, Nab2: interaction *F*_(2, 23)_= 3.343; p= 0.0532, Tukey post-test: *p<0.05. Egr3: interaction *F*_(2, 24)_= 19.20; p<0.0001, Tukey post-test: ***p<0.001, ****p<0.0001; Cut and Run chromatin n= 8-9 per group; Fig3 D). Consistent with the changes in the mRNA and protein levels of Nab2 and Egr3 when targeted with CRISPRi, H3K27ac enrichment on Nab2 and Egr3 promoters was decreased Nab2 sgRNA CRISPRi and Egr3 sgRNA CRISPRi targeting respectively (One-way ANOVA, Nab2: interaction *F*_(2, 22)_= 17.41; p< 0.0001, Tukey post-test: ***p<0.001, ****p<0.0001. Egr3: interaction *F*_(2, 24)_= 63.33; p<0.0001, Tukey post-test: ****p<0.0001; Cut and Run chromatin n= 7-9 per group; Fig3 D). These results demonstrate the target specificity of CRISPRi system and suggest that deacetylation at the H3K27 mark may drive the mRNA and protein level changes of target genes using the CRISPRi system. Consistent with our findings in CRISPRi Cut & Run experiments, the enrichment of CRISPRa complex on Nab2 and Egr3 promoter was increased in cells transfected with CRISPRa targeted with Nab2 and Egr3 sgRNAs, respectively, compared to lacZ sgRNA controls (One-way ANOVA, Nab2: interaction *F*_(2, 24)_= 9.147; p<0.01, Tukey post-test: **p<0.01. Egr3: interaction *F*_(2, 24)_= 11.71; p<0.001, Tukey post-test: **p<0.01, ***p<0.001; Cut and Run chromatin n= 9 per group; Fig3 E). Enrichment of H3K4me3 on the Nab2 and Egr3 promoters were significantly increased with Nab2 sgRNA CRISPRa and Egr3 sgRNA CRISPRa targeting respectively compared to lacZ sgRNA controls (One-way ANOVA, Nab2: interaction *F*_(2, 24)_= 13.14; p= 0.0001, Tukey post-test: ***p<0.001. Egr3: interaction *F*_(2, 24)_= 12.78; p<0.001, Tukey post-test: ***p<0.001; Cut and Run chromatin n= 9 per group; Fig3 E). Enrichment of H3K27ac on Nab2 promoter was trending toward a significant increase compared to lacZ control with Nab2 sgRNA CRISPRa targeting, and H3K27ac enrichment on Egr3 promoter was significantly increased relative to lacZ control with Egr3 sgRNA CRISPRa targeting (One-way ANOVA, Nab2: interaction *F*_(2, 24)_= 3.885; p= 0.0345, Tukey post-test: n.s.. Egr3: interaction *F*_(2, 24)_= 9.756; p<0.001, Tukey post-test: **p<0.01; Cut and Run chromatin n= 9 per group; Fig3 E). These results reveal the target specificity of the CRISPRa system, and suggest that H3K27 acetylation and H3K4 trimethylation may drive the mRNA and protein level changes of target genes using CRISPRa.

### Histone lysine demethylases display bidirectional induction in NAc D1-MSNs and D2-MSNs after repeated cocaine

Since EGR3 has a predicted binding sites with Kdm1a, Kdm6a, and Kdm5c (gene-regulation.com/pub/program/alibaba2) (Fig4 A), we next investigated NAc D1 and D2-MSN cell subtype ribosome-associated mRNA of Kdm1a, Kdm5c, and kdm6a, together with Kdm4a which does not have a putative binding site with EGR3, after seven daily injections of cocaine (20mg/kg) followed by a 24 hour abstinent period in D1-Cre-RT and D2-Cre-RT mice (Fig4 B). Interestingly, Kdm1a mRNA was increased in D1-MSNs, while it was decreased in D2-MSNs in the NAc of mice repeatedly exposed to cocaine compared to saline controls (Student’s *t* test, *p<0.05; mRNA sample n= 11 per group, *t*_(20)_= 2.279, p<0.05; mRNA sample n= 11 per group, *t*_(20)_= 2.279, p<0.05; Fig4 B). Similarly, Kdm6a mRNA was also increased in D1-MSNs, while decreased in D2-MSNs in this condition (Student’s *t* test, *p<0.05, ***p<0.001; mRNA sample n= 11 per group, *t*_(20)_= 2.436, p<0.05; mRNA sample n= 10-11 per group, *t*_(19)_= 4.364, p<0.001; Fig4 C). In contrast, Kdm5c mRNA was increased in D2-MSNs, while D1-MSNs did not show a significant change in Kdm5c mRNA expression with repeated exposure to cocaine (Student’s *t* test, **p<0.01; mRNA sample n= 11-12 per group, *t*_(21)_= 0.6308, p=0.5350; mRNA sample n= 11-12 per group, *t*_(21)_= 3.341, p<0.01; Fig4 D). Notably, we did not observe a change in Kdm4a, which does not have a putative EGR3 binding site, in either D1-MSNs or D2-MSNs after repeated exposure to cocaine (Student’s *t* test; mRNA sample n= 6 per group, *t*_(10)_= 0.4764, p=0.6440; mRNA sample n= 5 per group, *t*_(8)_= 2.060, p=0.0733; Fig4 E).

**Fig. 4.**
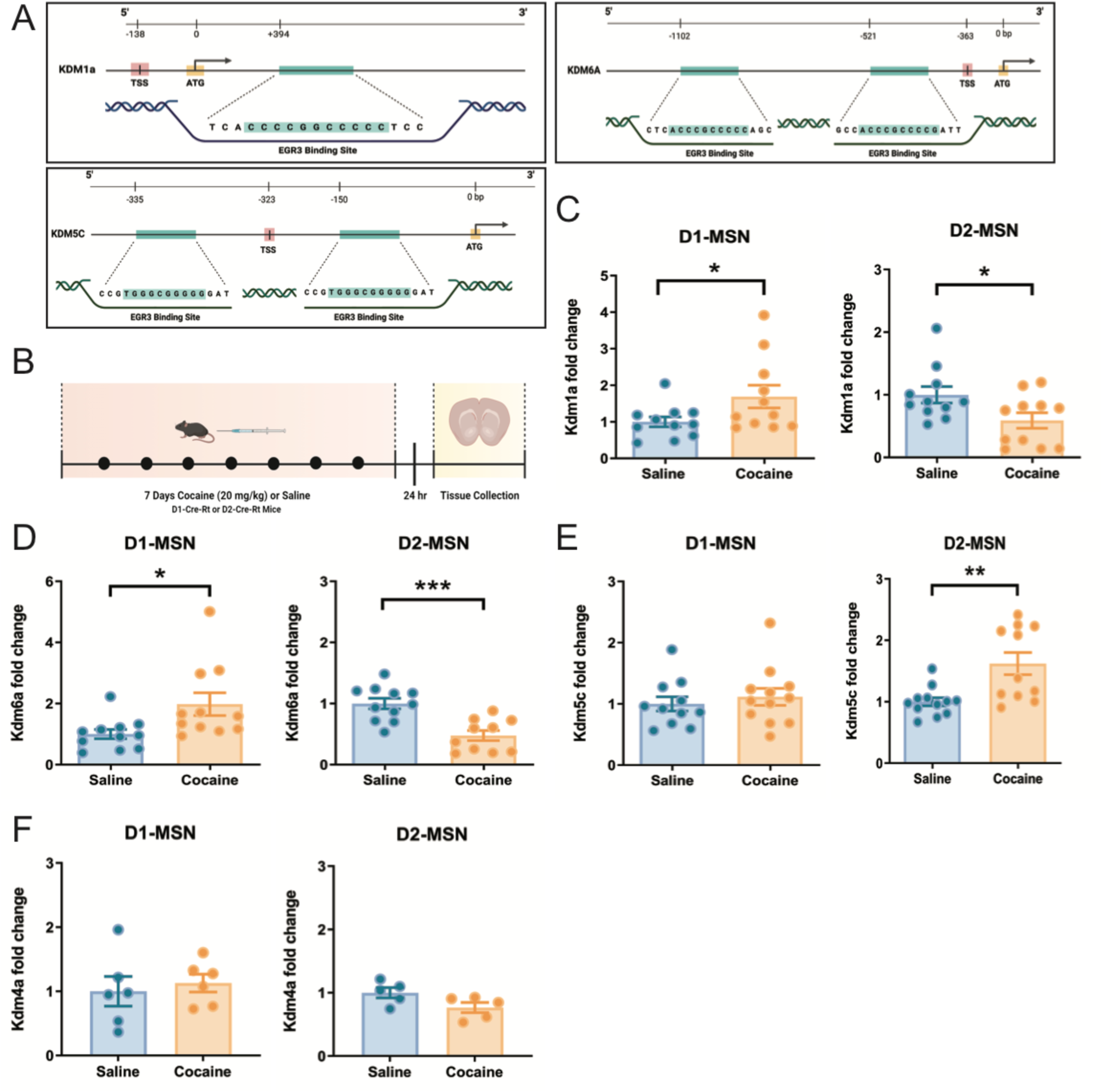
Kdms display bidirectional expression patterns in D1-MSNs and D2-MSNs after repeated cocaine exposure. **A**. Kdm1a, Kdm6a, and Kdm5c has GC rich EGR3 putative binding sites in the promoter and near the transcription start site. **B**. Male D1-Cre-RT and D2-Cre-RT mice received seven daily injections of cocaine (20mg/kg). Tissue punches were collected 24 hours post last injection. **C**. qRT-PCR shows increased Kdm1a mRNA in D1-MSNs while it decreased in D2-MSNs (D1-MSNs saline n= 11, cocaine n= 11; D2-MSNs saline n= 11, cocaine n= 11). **D**. Consistently, qRT-PCR shows increased Kdm6a mRNA levels in D1-MSNs while it decreased in D2-MSNs (D1-MSNs saline n= 11, cocaine n= 11; D2-MSNs saline n= 11, cocaine n= 10). **E**. qRT-PCR shows decreased Kdm5c mRNA levels in D2-MSNs while it did not change in D1-MSNs (D1-MSNs saline n= 11, cocaine n= 12; D2-MSNs saline n= 12, cocaine n= 11). **F**. No changes were observed in Kdm4a mRNA levels in either D1 or D2-MSNs (D1-MSNs saline n= 6, cocaine n= 6; D2-MSNs saline n= 5, cocaine n= 5). Each bar represents mean ± SEM. \**p*<0.05, \*\**p*<0.01, \*\*\**p*<0.001.

### Light inducible Opto-CRISPR-KDM1A and Opto-CRISPR-p300 system allows spatiotemporally precise perturbation of Nab2 and Egr3 expression

Given bidirectional induction of some KDMs in NAc cell subtypes after repeated cocaine, we aimed to explore if targeting KDM1A to Egr3 and Nab2 promoters can alter these transcripts in Neuro2A cells similar to cocaine exposure in NAc MSNs. To do this, we employed a Cre and light inducible CRISPR-KDM1A system (Opto-CRISPR-KDM1A) which utilizes the blue light inducible CRY2-CIB heterodimerizing system (Kennedy et al., 2010; Hughes et al., 2012; Konermann et al., 2013; Nihongaki et al., 2015; Polstein and Gersbach, 2015; Taslimi et al., 2016). By fusing the N-terminal fragment of CIB1 with dCas9 (DIO-dCas9-HA-CIBN), and truncated CRY2 with KDM1A (DIO-CRY2-FLAG-KDM1A), we developed a system to deliver KDM1A induced histone demethylation with spatiotemporal precision and Cre dependent specificity (Fig5 A). First, we transfected these vectors together with lacZ sgRNA in Neuro2a cells to confirm that these vectors express only in Cre positive cells. 48 hours post transfection, we immunostained cells for DAPI, GFP, HA-tag, and FLAG-tag. We observed GFP, HA-tag, and FLAG-tag signals only in Cre positive cells (Fig5 B). In Cre negative cells, we did not observe GFP, HA-tag, and FLAG-tag signals (Fig5 B). To address the possibility of functional promiscuity for unbound KDM1A, we compared the mRNA expression of Nab2 and Egr3 in Neuro2a cells transfected only with DIO-CRY2-FLAG-KDM1A and Cre vectors to cells transfected with Opto-CRISPR-KDM1A system targeted by lacZ sgRNA. In cells transfected only with DIO-CRY2-FLAG KDM1A and Cre vectors, we did not observe significant changes in Nab2 or Egr3 mRNA levels compared to cells transfected with Opto-CRISPR-KDM1A targeted by lacZ sgRNA after 2 hours of 1mW blue light stimulation (Student’s *t* test; mRNA sample n= 3 per group, *t*_(4)_= 0.2908, p=0.7857; mRNA sample n= 3 per group, *t*_(4)_= 0.8796, p=0.4287; Fig5 C). To investigate if blue light stimulation itself may alter the expression of our target genes, we compared the mRNA expression of Nab2 and Egr3 in Neuro2a cells transfected with Opto-CRISPR-KDM1A targeted by lacZ sgRNA in no light condition and 2 hours of 1mW blue light stimulation. Between these two conditions, we did not observe significant changes in Nab2 or Egr3 mRNA levels (Student’s *t* test; mRNA sample n= 3 per group, *t*_(4)_= 1.183, p=0.3023; mRNA sample n= 3 per group, *t*_(4)_= 2.126, p=0.1006; Fig5 D). To avoid potential blue light-induced gene expression alterations from phototoxicity, we changed the culture media to Photostable NEUMO media at least 3 hours prior to the start of light stimulations in all of the experiments performed in this study (Duke et al., 2019).

**Fig. 5.**
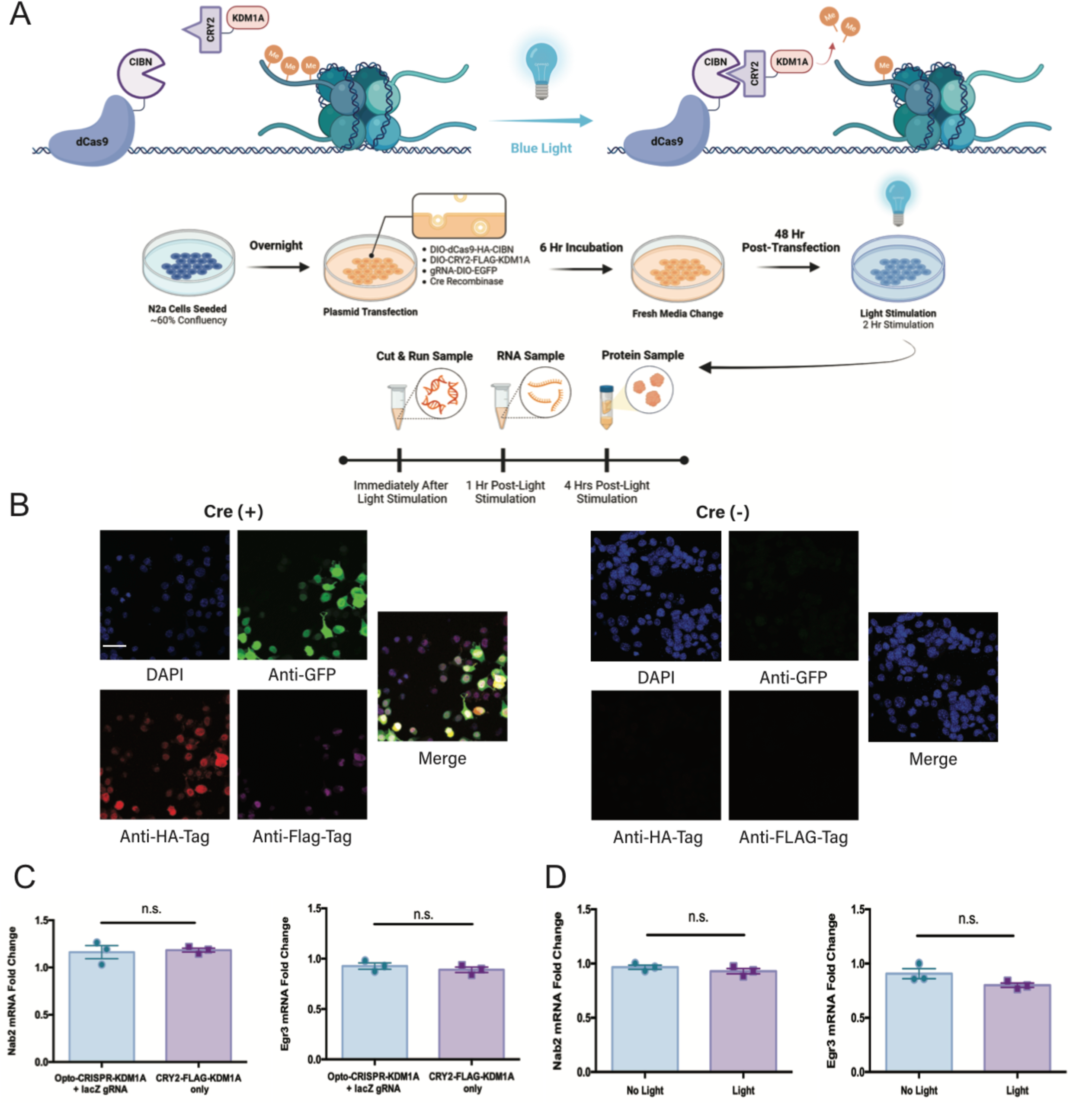
Development of a Cre and light inducible Opto-CRISPR-KDM1A system. **A**. Illustration of light inducible dCas9 and KDM1A fusion complex using CIBN and CRY2 heterodimers. Neuro2a cells received 2 hours of 1mW blue light stimulation. **B**. Immunocytochemistry of Neuro2a cells transfected with Opto-CRISPR-KDM1A display a Cre inducible expression as observed by GFP (green), HA-Tag (red), and Flag-Tag (magenta) expression. DAPI signal is blue. Scale bar, 50μM. **C**. qRT-PCR demonstrates cells transfected only with DIO-CRY2-FLAG-KDM1Aand Cre vectors do not show significant changes in Nab2 or Egr3 mRNA levels compared to cells transfected with complete Opto-CRISPR-KDM1A targeted by lacZ sgRNA after 2 hours of blue light stimulation (n= 3 per condition). **D**. qRT-PCR demonstrates that Neuro2a cells transfected with Opto-CRISPR-KDM1A with lacZ sgRNA in no light condition and 2 hours of 1mW blue light stimulation do not display significant differences in Nab2 and Egr3 mRNA levels (n= 3 per condition). Each bar represents mean ± SEM. n.s., not significant.

We next used the Opto-CRISPR-KDM1A with Nab2 sgRNA or Egr3 sgRNA to determine if recruiting KDM1A to these promoters can reduce or enhance transcription since KDM1A demethylates both positive and repressive histone marks. Using Opto-CRISPR-KDM1A with Nab2 sgRNA targeting, we observed a reduction in Nab2 mRNA expression compared to lacZ controls after 2 hours of 1mW blue light stimulation (One-way ANOVA, Nab2: interaction *F*_(3, 32)_= 145.8; p= 0.0001, Tukey post-test: **p<0.01, ****p<0.0001; mRNA n= 9 per group; Fig6 A). In Neuro2a cells transfected with Egr3 sgRNA and Opto-CRISPR-KDM1A, we also observed reduction of Egr3 mRNA expression compared to lacZ controls after 2 hours of 1mW blue light stimulation (One-way ANOVA, Egr3: interaction *F*_(3, 32)_= 84.44; p= 0.0001, Tukey post-test: ****p<0.0001; mRNA n= 9 per group; Fig6 A). Consistent with our findings with CRISPRi, Nab2 mRNA was upregulated compared to lacZ controls with Opto-CRISPR-KDM1A and Egr3 sgRNA targeting (One-way ANOVA, Nab2: interaction *F*_(3, 32)_= 145.8; p= 0.0001, Tukey post-test: **p<0.01, ****p<0.0001; mRNA n= 9 per group; Fig6 A). These results suggest that Opto-CRISPR-KDM1A interacts with nucleosome remodeling and histone deacetylase (NuRD) complex to cause demethylation at H3K4 methyl sites, resulting in transcriptional repression (Wang et al., 2009; Basta and Rauchman, 2015). Consistent observations were made at the protein level of NAB2 and EGR3 via western blots. 2 hours of 1mW blue light stimulation in Neuro2a cells transfected with Opto-CRISPR-KDM1A targeted by Nab2 sgRNA reduced NAB2 protein expression and induced EGR3 protein expression compared to lacZ controls (One-way ANOVA, NAB2: interaction *F*_(3, 8)_= 7.602; p= 0.01, Tukey post-test: *p<0.05. EGR3: interaction *F*_(3, 8)_= 47.89; p<0.0001, Tukey post-test: **p<0.01; Protein n= 3 per group; Fig6 B). In cells transfected with Opto-CRISPR-KDM1A targeted by Egr3 sgRNA, 2 hours of 1mW blue light stimulation reduced EGR3 protein expression compared to lacZ controls (One-way ANOVA, EGR3: interaction *F*_(3, 8)_= 47.89; p<0.0001, Tukey post-test: **p<0.01; Protein n= 3 per group; Fig6 B). To validate the specificity of Opto-CRISPR-KDM1A in our findings, we performed Cut & Run experiments with anti-HA to target the HA-tag on the dCas9-CIBN portion of the vectors, as well as antibodies against hallmark activation and repressive histone methylation and acetylation modifications H3K4me3 and H3K27ac (Shi et al., 2004; Wysocka et al., 2006; Robison and Nestler, 2011; Ferrari et al., 2014; Nestler, 2014). In Neuro2a cells transfected with Opto-CRISPR-KDM1A with Nab2 sgRNA, we observed an increase in the enrichment of Opto-CRISPR-KDM1A complex on Nab2 promoter compared to lacZ sgRNA controls (One-way ANOVA, Nab2: interaction *F*_(2, 24)_= 24.93; p<0.0001, Tukey post-test: ****p<0.0001; Cut and Run chromatin n= 9 per group; Fig6 C). In cells transfected with Opto-CRISPR-KDM1A with Egr3 sgRNA, we observed an increase in the enrichment of Opto-CRISPR-KDM1A complex on Egr3 promoter compared to lacZ sgRNA controls (One-way ANOVA, Egr3: interaction *F*_(2, 24)_= 16.01; p<0.0001, Tukey post-test: ***p<0.001; Cut and Run chromatin n= 9 per group; Fig6 C). Consistent with the changes in the mRNA and protein levels of Nab2 and Egr3 when targeted with Opto-CRISPR-KDM1A and their respective sgRNAs, H3K27ac and H3K4me3 enrichment on Nab2 promoter was decreased in Nab2 sgRNA Opto-CRISPR-KDM1A system and H3K27ac and H3K4me3 enrichment on Egr3 promoter was decreased in the Egr3 sgRNA Opto-CRISPR-KDM1A system (One-way ANOVA, H3K27ac Nab2: interaction *F*_(2, 24)_= 32.07; p<0.0001, Tukey post-test: **p<0.01, ***p<0.001, ****p<0.0001. H3K27ac Egr3: interaction *F*_(2, 24)_= 7.654; p<0.01, Tukey post-test: *p<0.05, **p<0.01. H3K4me3 Nab2: interaction *F*_(2, 23)_= 6.646; p<0.01, Tukey post-test: **p<0.01. H3K4me3 Egr3: interaction *F*_(2, 24)_= 5.188; p<0.05, Tukey post-test: *p<0.05; Cut and Run chromatin n= 8-9 per group; Fig6 C). These results demonstrate the target specificity of Opto-CRISPR-KDM1A system. Furthermore, these results show KDM1A targeting using our design of Nab2 and Egr3 sgRNA suggests a transcriptionally repressive H3K4 demethylase function for KDM1A rather than a transcription inducive H3K9 demethylase function (Shi et al., 2004; Metzger et al., 2005).

**Fig. 6.**
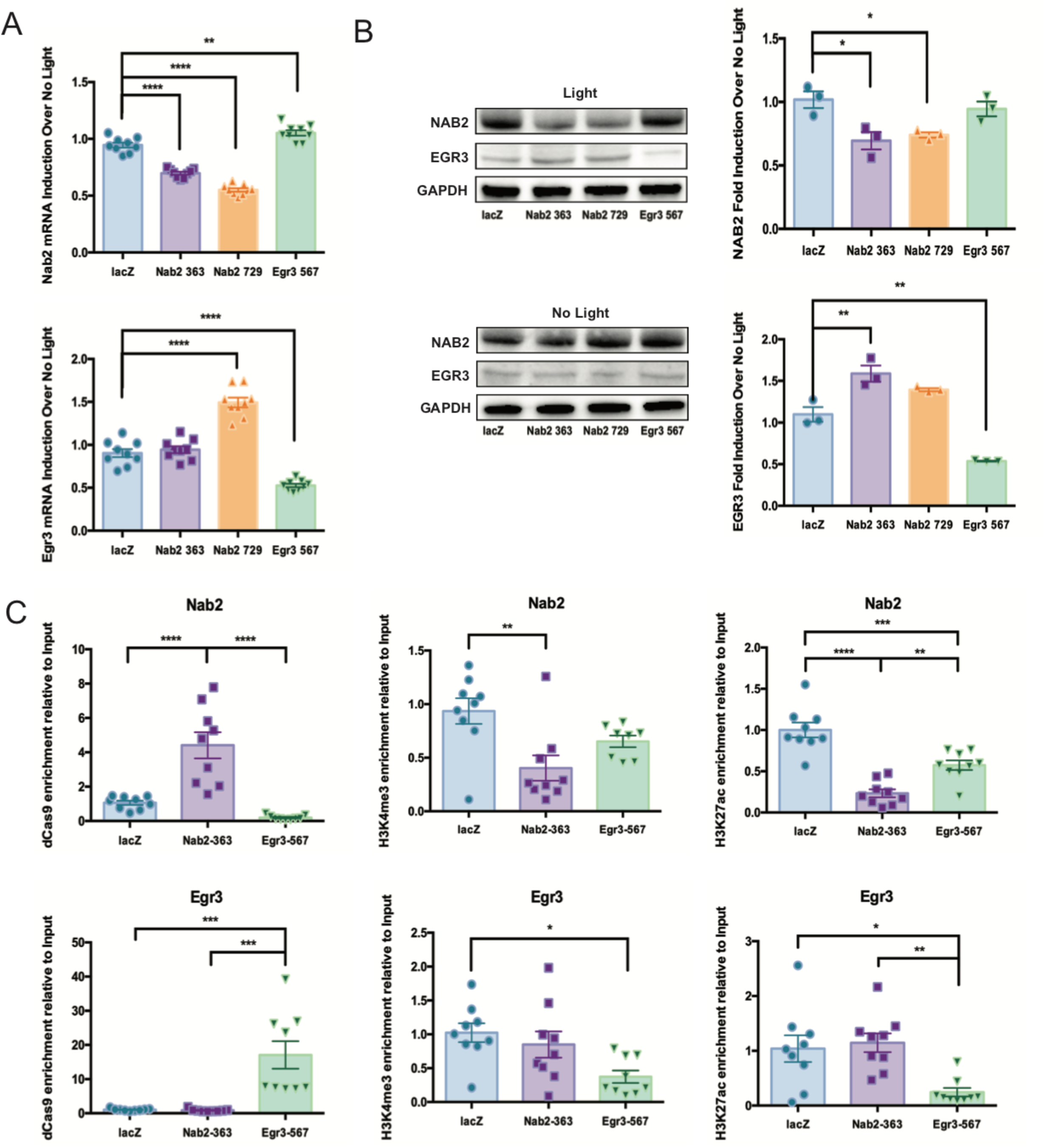
Opto-CRISPR-KDM1A mediated inhibition of Nab2 and Egr3 transcription. **A**. qRT-PCR demonstrates that 2 hours of blue light stimulation downregulated Nab2 mRNA while inducing Egr3 mRNA expression in cells transfected with Nab2 targeted Opto-CRISPR-KDM1A. Conversely, 2 hours of blue light stimulation induced Nab2 mRNA while downregulating Egr3 mRNA expression in cells transfected with Opto-CRISPR-KDM1A and Egr3 sgRNA (n= 9 per condition). **B**. Western blots show 2 hours of blue light stimulation downregulated NAB2 levels while upregulating EGR3 levels in cells transfected Opto-CRISPR-KDM1A and Nab2 sgRNA. EGR3 levels were downregulated while NAB2 levels were unchanged in cells transfected with Opto-CRISPR-KDM1A and Egr3 sgRNA (n= 3 per condition). **C**. Cut and Run on blue light stimulated Neuro2a cells transfected with Opto-CRISPR-KDM1A shows enrichment of Opto-CRISPR-KDM1A complex on the promoters of respective genes targeted by sgRNAs compared to lacZ sgRNA controls. Transcriptional activation marks, H3K4me3 and H3K27ac, have reduced enrichment on the promoters of respective genes targeted by sgRNAs compared to lacZ sgRNA controls (n= 9 per condition). Each bar represents mean ± SEM. \**p*<0.05, \*\**p*<0.01, \*\*\**p*<0.001, \*\*\*\**p*<0.0001.

Since Opto-CRISPR-KDM1A acts to repress transcription, Opto-CRISPRi, we then wanted to use a Opto-CRISPRa approach. To do this we developed a new opto-CRISPR-p300 vector in which KDM1A portion of DIO-CRY2-FLAG-KDM1A vector was replaced by the truncated p300 functional core (DIO-CRY2-FLAG-p300Core) to induce histone acetylation at the H3K27 mark (Fig7 A) (Delvecchio et al., 2013; Hilton et al., 2015). We first performed immunohistochemistry in Neuro2a cells transfected with Opto-CRISPR-p300, lacZ sgRNA and Cre to demonstrate that the vectors express only within Cre positive cells (Fig7 B). To address the possibility of functional promiscuity for unbound p300core, we compared the mRNA expression of Nab2 and Egr3 in Neuro2a cells transfected only with DIO-CRY2-FLAG-p300core and Cre vectors to cells transfected with Opto-CRISPR-p300 and lacZ sgRNA. In cells transfected only with DIO-CRY2-FLAG-p300core, we did not observe significant changes in Nab2 or Egr3 mRNA levels compared to cells transfected with Opto-CRISPR-p300 and lacZ sgRNA after 2 hours of 1mW blue light stimulation (Student’s *t* test; mRNA sample n= 3 per group, *t*_(4)_= 0.7803, p=0.4788; mRNA sample n= 3 per group, *t*_(4)_= 0.6398, p=0.5571; Fig7 C). In determine if blue light stimulation itself may alter the expression of our target genes, we compared the mRNA expression of Nab2 and Egr3 in Neuro2a cells transfected with Opto-CRISPR-p300 with lacZ sgRNA in no light condition and 2 hours of 1mW blue light stimulation. Between these two conditions, we did not observe significant changes in Nab2 or Egr3 mRNA levels (Student’s *t* test; mRNA sample n= 3 per group, *t*_(4)_= 2.072, p=0.1070; mRNA sample n= 3 per group, *t*_(4)_= 1.098, p=0.3340; Fig7 D)..

**Fig. 7.**
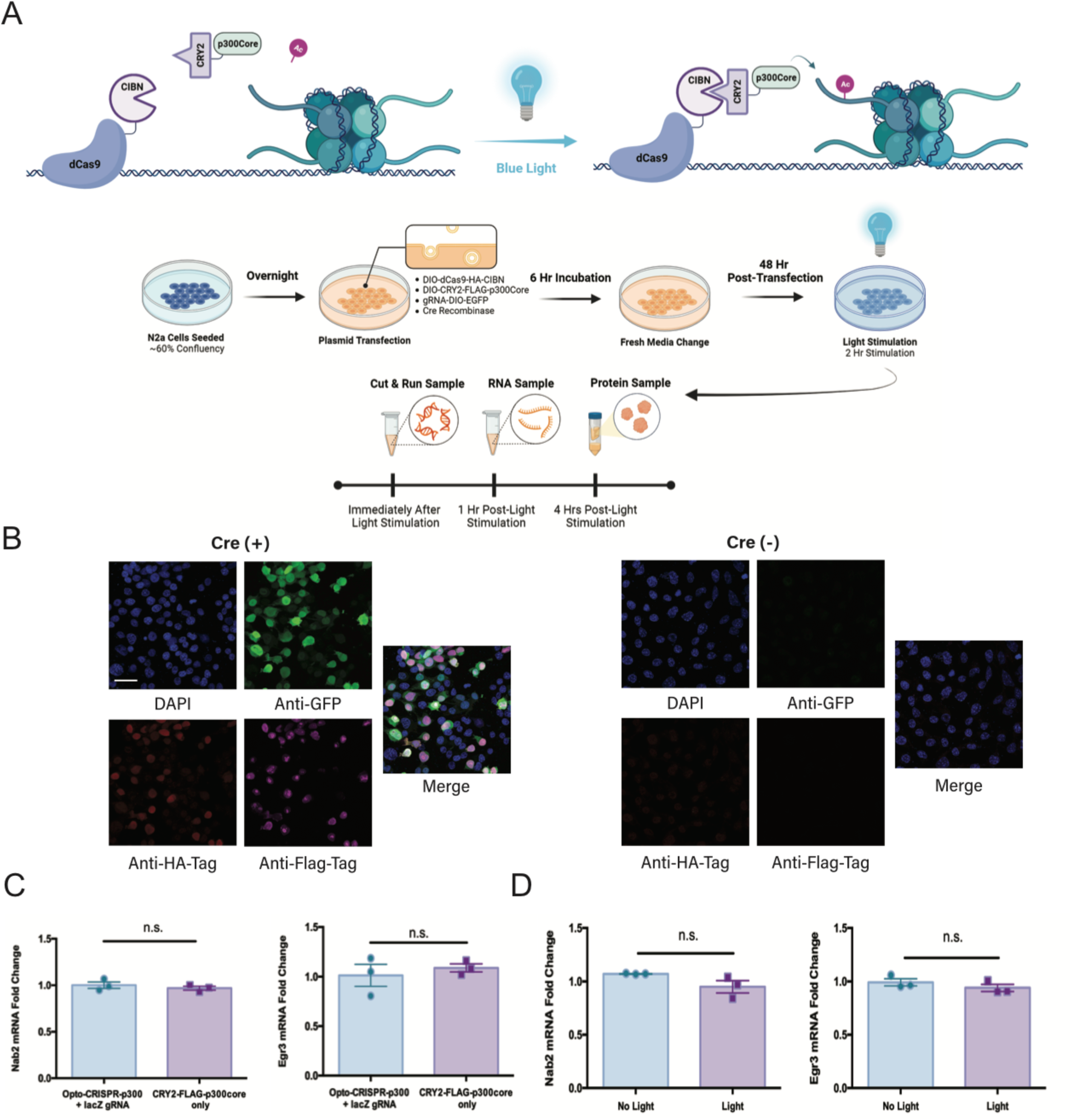
Development of a Cre and light inducible Opto-CRISPR-p300 system. **A**. Illustration of light inducible dCas9 and p300 fusion complex using CIBN and CRY2 heterodimers. Neuro2a cells received 2 hours of 1mW blue light stimulation. **B**. Immunocytochemistry of Neuro2a cells transfected with Opto-CRISPR-p300 system demonstrate a Cre inducible expression as observed by GFP (green), HA-Tag (red), and Flag-Tag (magenta) expression. DAPI signal is blue. Scale bar, 50μM. **C**. qRT-PCR demonstrates that the expression of DIO-CRY2-FLAG-p300core alone does not induce changes in Nab2 or Egr3 mRNA levels compared to cells transfected with Opto-CRISPR-p300 targeted by lacZ sgRNA after 2 hours of blue light stimulation (n= 3 per condition). **D**. qRT-PCR demonstrates Neuro2a cells transfected with Opto-CRISPR-p300 targeted by lacZ sgRNA in no light condition and 2 hours of 1mW blue light stimulation do not display significant differences in Nab2 and Egr3 mRNA levels (n= 3 per condition). Each bar represents mean ± SEM. n.s., not significant.

We then transfected Opto-CRISPR-p300 with Nab2 sgRNA or Egr3 sgRNA into Neuro2a cells. After 2 hours of 1mW blue light stimulation, Nab2 mRNA and Egr3 mRNA were significantly increase compared to lacZ controls when targeting with their respective gRNAs (One-way ANOVA, Nab2: interaction *F*_(3, 32)_= 3.872; p<0.05, Tukey post-test: *p<0.05. Egr3: interaction *F*_(3, 32)_= 13.03; p<0.0001, Tukey post-test: **p<0.01; mRNA n= 9 per group; Fig8 A). These results were consistent in protein expression changes demonstrated via western blots, as 2 hours of 1mW blue light stimulation in Neuro2a cells transfected with Opto-CRISPR-p300 with Nab2 sgRNA induced NAB2 protein expression. Notably, EGR3 expression was induced in these cells compared to lacZ controls, although this bidirectional regulation was not observed at the mRNA level (One-way ANOVA, NAB2: interaction *F*_(3, 8)_= 5.296; p<0.05, Tukey post-test: *p<0.05. EGR3: interaction *F*_(3, 8)_= 19.78; p<0.001, Tukey post-test: *p<0.05; Protein n= 3 per group; Fig8 B). In cells transfected with Opto-CRISPR-p300 targeted by Egr3 sgRNA, 2 hours of 1mW blue light stimulation induced EGR3 protein expression compared to lacZ controls (One-way ANOVA, EGR3: interaction *F*_(3, 8)_= 19.78; p<0.001, Tukey post-test: *p<0.05; Protein n= 3 per group; Fig8 B). To validate the specificity of Opto-CRISPR-p300, we performed Cut & Run experiments with anti-HA to target the HA-tag on the dCas9-CIBN as well as anti-H3K4me3 and anti-H3K27ac. In Neuro2a cells transfected with Opto-CRISPR-p300 with Nab2 sgRNA, we saw an increase in the enrichment of Opto-CRISPR-p300 complex on the Nab2 promoter compared to lacZ sgRNA controls (One-way ANOVA, Nab2: interaction *F*_(2, 23)_= 35.67; p<0.0001, Tukey post-test: ****p<0.0001; Cut and Run chromatin n= 8-9 per group; Fig8 C). When using the Egr3 sgRNA instead, we observed an increase in the enrichment of Opto-CRISPR-p300 complex on the Egr3 promoter compared to lacZ sgRNA controls (One-way ANOVA, Egr3: interaction *F*_(2, 24)_= 10.31; p<0.001, Tukey post-test: **p<0.01; Cut and Run chromatin n= 9 per group; Fig8 C). Notably, H3K4me3 enrichment on the Nab2 promoter was not significantly changed in the Nab2 sgRNA Opto-CRISPR-p300 system, while H3K4me3 enrichment on the Egr3 promoter was decreased with Egr3 sgRNA and Opto-CRISPR-p300 (One-way ANOVA, Nab2: interaction *F*_(2, 22)_= 4.291; p<0.05, Tukey post-test: *p<0.05. Egr3: interaction *F*_(2, 24)_= 17.82; p<0.0001, Tukey post-test: ***p<0.001 ****p<0.0001; Cut and Run chromatin n= 7-9 per group; Fig8 C). Consistent with the changes in the mRNA and protein levels of Nab2 and Egr3 when targeted with Opto-CRISPR-p300 and their respective sgRNAs, H3K27ac enrichment on the Nab2 promoter was increased in Nab2 sgRNA Opto-CRISPR-p300 system and H3K27ac enrichment on Egr3 promoter was increased in the Egr3 sgRNA Opto-CRISPR-p300 system (One-way ANOVA, Nab2: interaction *F*_(2, 24)_= 8.603; p<0.01, Tukey post-test: **p<0.01. Egr3: interaction *F*_(2, 23)_= 20.14; p<0.0001, Tukey post-test: ****p<0.0001; Cut and Run chromatin n= 9 per group; Fig8 C). These results demonstrate the target specificity of the Opto-CRISPR-p300 system. Furthermore, these results show p300core targeting using our Opto-CRISPR-p300 system induces activation of gene expression mainly driven by acetylation of H3K27 mark.

**Fig. 8.**
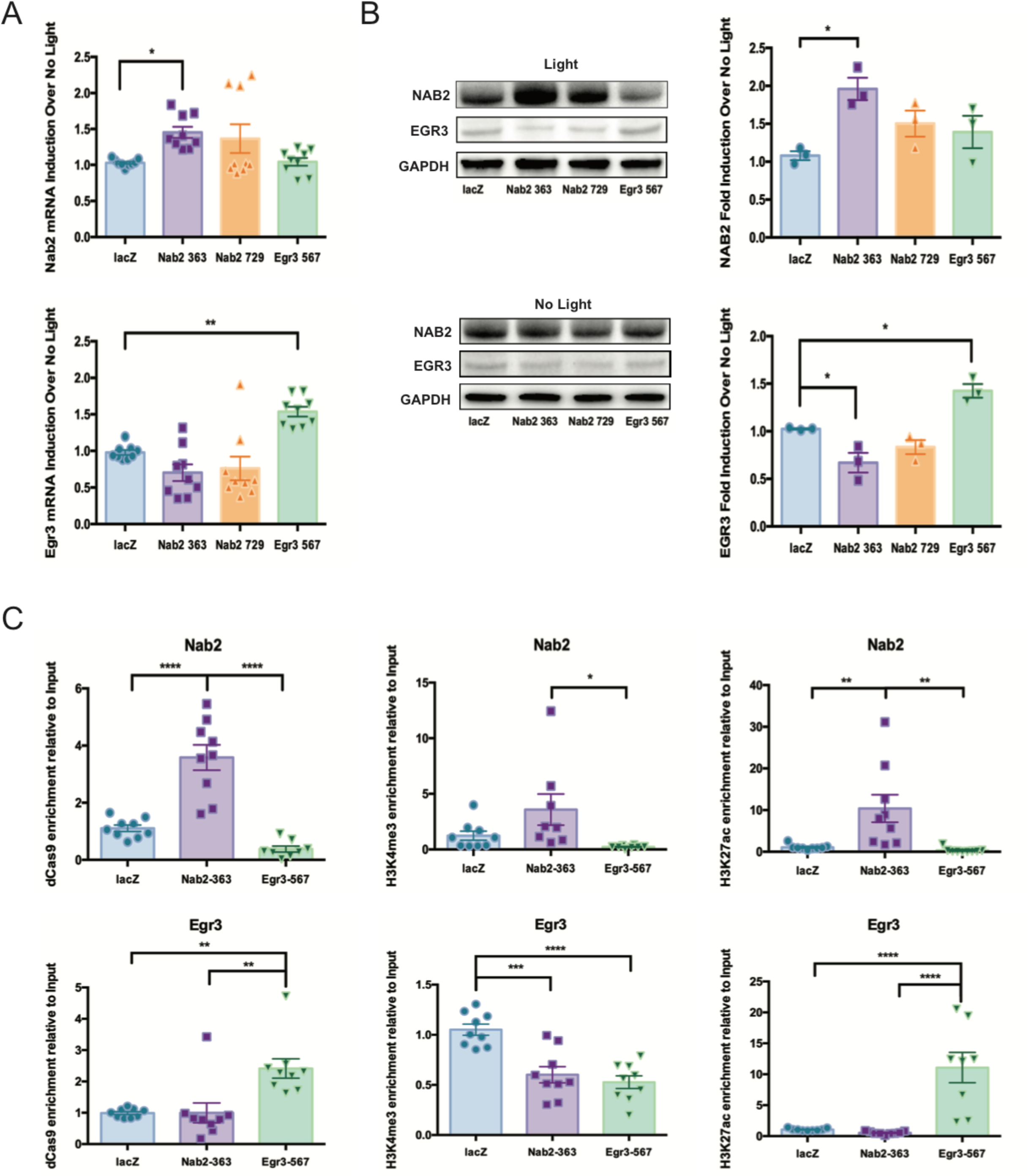
Opto-CRISPR-p300 mediated activation of Nab2 and Egr3 transcription. **A**. qRT-PCR demonstrate that 2 hours of blue light stimulation induced Nab2 mRNA and Egr3 mRNA expressios in cells transfected with Nab2 or Egr3 sgRNAs respectively and Opto-CRISPR-p300 (n= 9 per condition). **B**. Western blots demonstrate that 2 hours of blue light stimulation induced NAB2 levels while downregulating EGR3 levels in cells transfected with Opto-CRISPR-p300 and Nab2 sgRNA. EGR3 levels were upregulated while NAB2 levels were unchanged in cells transfected with Opto-CRISPR-p300 and Egr3 sgRNA (n= 3 per condition). **C**. Cut and Run of blue light stimulated Neuro2a cells transfected with Opto-CRISPR-p300 display enrichment of Opto-CRISPR-p300 complex on the promoters of respective genes targeted by sgRNAs compared to lacZ sgRNA controls. Transcriptional activation marks, H3K4me3 and H3K27ac, have increased enrichment on the promoters of respective genes targeted by sgRNAs compared to lacZ sgRNA controls (n= 9 per condition). Each bar represents mean ± SEM. \**p*<0.05, \*\**p*<0.01, \*\*\**p*<0.001, \*\*\*\**p*<0.0001.

## Discussion

Our study demonstrates bidirectional regulation of Nab2 in NAc MSN subtypes after repeated cocaine exposure. Nab2 was reduced in D1-MSNs and enhanced in D2-MSNs. As our previous work showed Egr3 is reduced in D2-MSNs after repeated cocaine (Chandra et al., 2015), the current data is consistent with Nab2 acting in concert with Egr3 to reduce Egr3 expression (Kumbrink et al., 2010). Not surprisingly in D1-MSNs, when Egr3 is induced, after repeated cocaine exposure, we observed a reduction of its co-transcriptional repressor, Nab2. This intricate regulation was masked in bulk tissue where Nab2 mRNA was not significantly changed in mice repeatedly exposed to cocaine. Although our previous study demonstrates a modest reduction of Egr3 in bulk NAc tissue in this condition (Chandra et al., 2015). These findings emphasize the possibility of missing critical molecular adaptations when the observations are made at the whole tissue level (Chandra et al., 2015).

Previous findings have shown that Nab2 is induced via BDNF signaling (Chandwani et al., 2013). Furthermore, previous reports have demonstrated that Egr3 is regulated via BDNF signaling (Roberts et al., 2006; Kim et al., 2012). BDNF may act preferentially in D2-MSNs through higher expression of BDNF and TrkB receptors, which may induce Egr3 and Nab2 (Lobo et al., 2010). This may drive a negative feedback loop in which Egr3 self-regulates its own transcription via induction of its corepressor Nab2. In contrast, the Egr3 increase in D1-MSNs may occur through its positive feedback loop by which Egr3 binds to its own promoter to enhance Egr3 transcription (Kumbrink et al., 2010). D1 and D2 receptor signaling may also contribute to this cocaine driven bidirectional regulation of Nab2 and Egr3 in D1 and D2-MSNs. Previous findings have shown that D1 receptor signaling could potentially enhance cAMP-PKA-CREB signaling cascades (Carlezon et al., 2005; Surmeier et al., 2007; Suehiro et al., 2010). On the contrary, Gi coupled D2 receptor signaling may inhibit cAMP-PKA-CREB signaling cascades (Carlezon et al., 2005; Surmeier et al., 2007; Suehiro et al., 2010). We speculate that the changes occurring downstream of these signaling cascades may induce Egr3 in D1-MSNs, which will ultimately drive Nab2 levels down in a negative feedback loop to reduce Egr3 in D2-MSNs while Nab2 levels remain elevated.

Consistent with the cocaine driven bidirectional regulation of Nab2 and Egr3, we observed an increase in Egr3 when Nab2 was silenced using CRISPRi in Neuro2a cells. This feedback mechanism was also observed by an increase in Nab2 when blocking Egr3 transcription. On the contrary, we observed reduced Egr3 levels when Nab2 was induced using CRISPRa in Neuro2a cells. When we enhanced Egr3, we observed reduced Nab2 levels. These bidirectional regulations closely mimicked cocaine driven regulation of Nab2 and Egr3 observed in D1- and D2-MSNs. Previous studies from our group have shown D1-MSN specific down-regulation of Egr3 and D2-MSN specific induction of Egr3 successfully blunted cocaine driven behaviors (Chandra et al., 2015; Engeln et al., 2020). Our CRISPR tools which express in Cre dependent manner afford the ability to manipulate MSN subtypes for *in vivo* gene perturbation studies to elucidate if D1-or D2-MSN Egr3 and Nab2 dynamics in the intact brain in the future studies.

Previous studies demonstrated altered histone methylation, and corresponding histone modifying enzymes, in NAc and other reward related brain regions with exposure to psychostimulants (Maze et al., 2010; Robison and Nestler, 2011; Aguilar-Valles et al., 2014; Heller et al., 2016). Genetic and pharmacological interventions of histone modifying enzymes in NAc and other reward related brain regions during drug mediated behaviors have shed light on epigenetic regulations and corresponding gene regulations. However, many of these studies examine global regulation of histone modifying enzymes rather than regulation at specific loci, although some recent studies examine loci specific manipulations (Carpenter et al., 2020; Xu et al., 2021). Furthermore, many of these studies examine bulk tissue alterations rather than cell subtype specific changes. In our findings, we observed bidirectional changes in expression levels for Kdm1a, and Kdm6a in D1 and D2-MSNs after repeated exposure to cocaine. In contrast, Kdm5c was only induced in D2-MSNs, while Kdm4a did not show a change in either subtype. The expression of Kdm6a in the prefrontal cortex was reported to be downregulated in rat cocaine self-administration studies (Sadakierska-Chudy et al., 2017). These changes in Kdms may occur through cocaine induced dopamine receptor signaling cascades. Previous studies have shown cocaine mediated D1 activation may cause imbalance of H3K4me3/H3K27me3 via downregulation of Kdm1a (González et al., 2020). These cell subtype specific regulation of Kdms may drive the imbalance of D1- and D2-MSNs epigenome regulation with repeated cocaine exposure, with impact on the transcriptional landscape to promote cellular and behavioral occurring with cocaine exposure.

Our findings that show bidirectional regulation of Kdm1a in NAc cell subtypes after repeated cocaine exposure emphasize the importance of examining Kdm1a regulation of Egr3 and Nab2 transcription. Using our Cre and blue light inducible Opto-CRISPR-KDM1A system to target KDM1A to the Nab2 promoter we observed reduced expression of Nab2 while inducing Egr3 expression in Neuro2a cells. Similarly, targeting the Egr3 promoter resulted in reduced expression of Egr3 while inducing Nab2 expression. Previous studies demonstrated that Kdm1a acts on H3K4 and H3K9 to demethylate these histone modifications (Shi et al., 2004; Forneris et al., 2005; Metzger et al., 2005; Garcia-Bassets et al., 2007; Nair et al., 2010; Rudolph et al., 2013). Kdm1a can function to both repress and activate transcription by mediating histone H3K4me1/2 and H3K9me1/2 demethylation respectively. Our findings show that Opto-CRISPR-KDM1A causes repression of both Egr3 and Nab2 and likely targets H3K4 at their promotors. However, it is plausible that sgRNA targeting at different promoter regions or other genes, could result in Opto-CRISPR-KDM1A demethylating H3K9 to induce transcription. Replacing the KDM1A with truncated core of p300 histone acetyltransferase allowed us to engineer a Opto-CRISPR-p300 system for Cre and light inducible activation of target genes. Targeting Nab2 with Opto-CRISPR-p300 induced the expression of Nab2 while reducing Egr3. In contrast targeting Egr3 with Opto-CRISPR-p300 induced the expression of Egr3 and blunted Nab2 expression in Neuro2a cells. This is consistent with previous findings which demonstrate that dCas9 fused catalytic core region of p300 is sufficient to drive the transcription of its target (Hilton et al., 2015) and consistent with the role of p300 targeting acetylation to promote transcription. Acetylation of H3K27 mark is one of the most widely studied indicators of enhancer activity (Ogryzko et al., 1996; Chen and Li, 2011; Delvecchio et al., 2013). Not surprisingly enrichment of H3K27ac on the promoters of Egr3 and Nab2, when Opto-CRISPR-p300 is targeted with Egr3 or Nab2 sgRNA respectively, is also consistent with endogenous p300 binding profiles (Visel et al., 2009; Rada-Iglesias et al., 2011).

In recent years, CRISPR tools have emerged as a potent method for genetic perturbation and epigenetic editing (Dominguez et al., 2016; Yeo et al., 2018; Zheng et al., 2018; Savell et al., 2019; Duke et al., 2020; Carullo et al., 2021). These tools offer considerable merit over the traditional methods for gene silencing and overexpression in sequence dependent and sequence independent off-target specificity (Bridge et al., 2003; Jackson et al., 2003; Sledz et al., 2003; Khan et al., 2009; Olejniczak et al., 2016). Since dCas9-KRAB and dCas9-VP64 mediated genetic perturbations can be achieved in Cre inducible cell subtype specific manner, these tools would have a broad appeal for studies that aim to decipher cell subtype specific questions. Our Opto-CRISPR tools allow Cre and light inducible spatio-temporally precise manipulations which may mimic more endogenous expression patterns and allow precise control of the duration of gene perturbation as well as reversible manipulations once the light stimulation has been turned off. *In vivo* applications of these Opto-CRISPR tools would allow temporal studies where perturbation of a target gene may be turned on at different time points during the course of the drug exposure.

Our findings show novel and dynamic regulation of Nab2 and Egr3 in NAc MSN subtypes and demonstrate the ability to use CRISPR epigenome editing tools to explore this regulation. The synthetic epigenetic manipulation of histone profiles and downstream transcription is critical in deciphering complex psychostimulant induced changes in molecular landscape. The versatility of these Cre and light inducible Opto-CRISPR tool sets allow the study of specific cell type transcriptional profiles after cocaine and other conditions. Importantly such tools can be applied in future studies for specific manipulation of transcriptional profiles that are critical for psychostimulant-induced cellular and behavioral plasticity and with spatiotemporal precision. These results emphasize the need for detailed elucidation of transcriptional underpinnings at cell subtype specific resolution and targeting cell subtype specific delivery of CRISPRi/a constructs allow physiologically relevant manipulations of transcripts to explore molecular substrates occurring in models of cocaine use disorder.

## Acknowledgements

We thank A. Poulopoulos, R. Richardson, T.A. Blanpied, and A.D. Levy (University of Maryland School of Medicine) for the advice and shared resources for Light Plate Apparatus experiments.

